# Not too rigid or too wobbly: defining an optimal membrane fluidity range essential for biofilm formation in *Escherichia coli*

**DOI:** 10.1101/2025.02.02.635587

**Authors:** Yaoqin Hong, Jilong Qin, Makrina Totsika

**Author notes:** For correspondence, contact Yaoqin Hong and Makrina Totsika.

## Abstract

**IMPACT STATEMENT:** Membrane fluidity plays a crucial role in bacterial fitness and adaptation to cope rapid environmental changes. While high membrane fluidity promotes robust biofilm formation in *Klebsiella pneumoniae*, studies in several other species, including *Salmonella enterica*, suggest that biofilm formation is associated with reduced fluidity. This paradox may reflect the complex relationship between lipid composition and biofilm formation. Our findings demonstrated that both low and high extremes of lipid fluidity restrict biofilm formation. We propose that the required fluidity for biofilm growth, relative to that required for planktonic growth, may differ between species and are readily adjusted during lifestyle transitions.

## CORRESPONDENCE

Microbes in natural environments often exist in communities, called biofilms, where cells are encased in an extracellular polymeric matrix (EPS) that provides protection from external stresses such as detergents and antibiotics. This EPS, is primarily composed of polysaccharides, proteins, lipids, and extracellular DNA, with its composition and spatial organization tightly regulated by molecular signals ^1^. Identifying the molecular cues that bacteria use to regulate biofilm formation and dispersal is critical for both combating infections and harnessing beneficial biofilms. The contribution of bacterial cell membrane fluidity, as a modulator of these processes, remains an underexplored area.

The bacterial membrane plays a vital role in supporting essential processes, including energy generation, fitness in different environments and changes in environmental conditions necessitates membrane lipid adjustments for survival and fitness ^2^. Changes in membrane lipids serve as a molecular cue in the Envelope Stress Response that can influence various pathways involved in biofilm development. A typical bacterial phospholipid contains two fatty acid (FA) chains, and the chemical nature of these FAs dictates the membrane fluidity ^2^. Previous studies have demonstrated that biofilms of *Salmonella enterica, Staphylococcus aureus, Listeria monocytogenes*, and *Pseudomonas aeruginosa* contain elevated levels of saturated fatty acids (SFAs) compared to their planktonic counterparts ^3^. This suggests that bacterial membrane composition and fluidity may play roles in biofilm formation, although the underlying mechanisms remain unclear ^3,4^.

Most bacterial species *de novo* synthesise the acyl chains to be integrate into membrane lipids by the highly conserved type II fatty acid synthesis pathway distinctive from that used by eukaryotes ^2^. As such, the pathway had served as an important target for antimicrobial discovery ^5,6^, with the recent progress shown the potential of potentiating susceptibility to otherwise ineffective antibiotics ^7,8^. In *Escherichia* coli, the major unsaturated fatty acids (UFAs) are palmitoleate (C16:1) and cis-vaccenate (C18:1). Bacterial phospholipids exhibit chemical asymmetry, with SFAs typically occupying the *sn*1 position and unsaturated fatty acids (UFAs) occupying the *sn*2 position of the glycerol backbone owed to the substrate preference of acyltransferases involved in phosphatidic acid synthesis ^2,9^. The UFA/SFA ratio that affects membrane fluidity is predominantly determined by the diversity of FAs at the *sn*2 position ^2,9^. The only exception to this substrate selectivity at position *sn*1 is the integration of cis-vaccenate, which leads to higher membrane fluidity that is imperative for bacterial fitness under certain conditions, such as low temperature growth ^10^. As such, this phospholipid asymmetry plays a pivotal role in modulating membrane fluidity, which, in turn, may influences biofilm formation. In *E. coli*, UFA synthesis is mediated by ketoacyl-ACP synthase I (*FabB*), while the conversion of palmitoleate to *cis*-vaccenate requires ketoacyl-ACP synthase II (*FabF*) ^11,12^. Thus, *FabB* and, to a lesser extent, *FabF*, are key players governing membrane fluidity. Interestingly, while *FabB* catalyses the rate limiting step in UFA synthesis and governs the fundamental basis of membrane fluidity, the enzyme is not thermoregulated ^13^. Rather, the *FabF* enzyme involved in the final conversion from palmitoleate to cis-vaccenate was reported to be under thermal regulation across phyla, including the Firmicutes ^14^. The two enzymes function concertedly to regulate membrane fluidity and optimise fitness required for survival across a wide range of temperatures.

The *E. coli* K-12 strain AR3110 exhibits a distinct macrocolony architecture when grown on agar surfaces, characterized by a radial, rugose, three-dimensional structure with delaminated buckling and skin-like elasticity ^15^. This strain also forms a robust pellicle highly resistant to mechanical disturbance at the liquid-air interface (Fig. 1B). Remarkably, both the rugose colony architecture and pellicle formation disappeared in the Δ*fabF* strain, suggesting that reduced membrane fluidity underlies these phenotypic changes (Fig. 1A and 1B).

**Fig 1.**
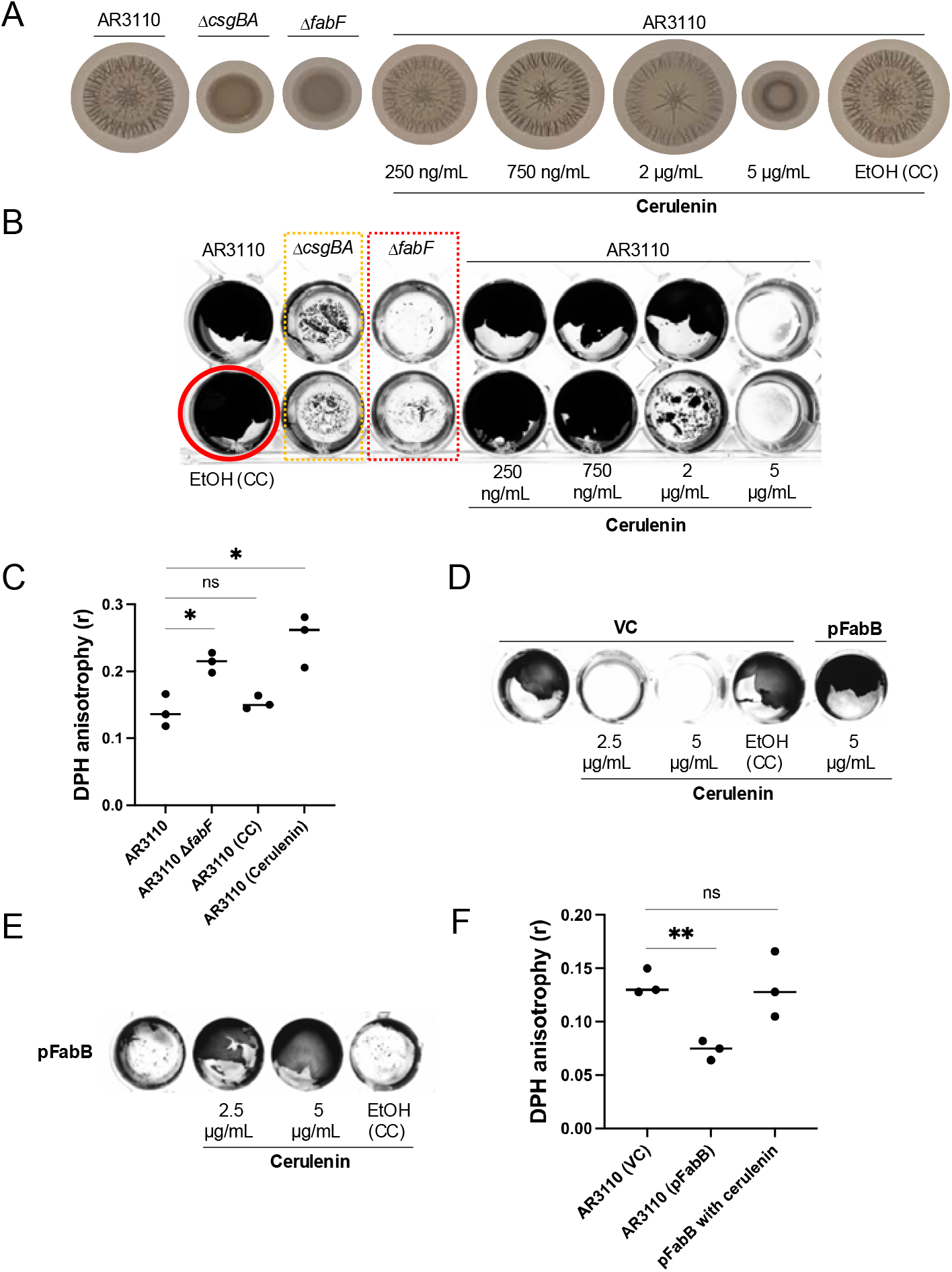
Optimal membrane fluidity governs *E. coli* biofilm rugosity and pellicle formation. (A) The *fabF* null mutation or cerulenin treatment eliminated rugosity in AR3110 macrocolony biofilms grown on YESCA agar at 26°C; (B) The *fabF* null mutation or cerulenin treatment abolished pellicle formation in AR3110 cultured in Mueller Hinton broth. Identical results were observed on YESCA, LB Lennox, and LBON; (C) DPH fluorescence anisotropy was used to measure membrane fluidity (note: fluidity is inversely proportional to fluorescence anisotropy); (D) Ectopic expression of *fabB* (pSU2718 vector, induced with 250μM IPTG) restored pellicle formation inhibited by cerulenin treatment (E) Overexpression of *fabB* blocked pellicle formation in AR3110, but this defect was alleviated by cerulenin treatment. (F) Ectopic expression of *FabB* significantly increased membrane fluidity; this increase was ameliorated by cerulenin treatment. AR3110 ΔcsgBA strain used in panel A is unable to synthesise curli ^15^, an essential structural component of AR3110 rugose macrocolonies. Panels A, B, D and E are representative of at least three independent observations. Data in Panels C and F were obtained from three independent repeats. Error bars indicate standard deviation. Statistical significance was determined using an unpaired, two-tailed Student’s t-test. P□<□0.05 was considered statistically significant. Asterisks indicate significance levels: ns = not significant (P□≥□0.05); P□<□0.05 (*); P□<□0.01 (**). Abbreviations: CC, carrier control (EtOH, ethanol used to dissolve cerulenin); VC, vector control (pSU2718).

Aside from the SFA/UFA ratio, membrane fluidity is also influenced by the profile of phospholipid headgroups. For example, phosphatidylethanolamine has a smaller headgroup than phosphatidylcholine, in this case, allowing formation of hydrogen bonds between cationic amino headgroups and the anionic phosphate residue of nearby lipids to affect membrane packing ^16,17^. Consistent with this, a reduction of membrane phosphatidylethanolamine content was observed for yeast cells at higher temperature growth ^18^. Weibel and colleagues also demonstrated cardiolipin deficiency results in the reduction of biofilm formation in *E. coli* ^4^. In this work, rather than using standard FA profiling, we utilised DPH fluorescence anisotropy, a common method for measuring membrane fluidity. A reduced anisotropy correlates with higher membrane fluidity, while higher anisotropy indicates a more ordered and less fluid membrane. The measured anisotropy in the Δ*fabF* strain was significantly higher compared to the wild-type AR3110 control (*t*-test, *P*=0.0111) (Fig. 1C), which supported the hypothesis that membrane fluidity is crucial for biofilm formation in *E. coli* (Fig. 1C).

Cerulenin inhibits both FabF and FabB, enzymes crucial for the synthesis of UFAs, thereby reducing membrane fluidity ^19^. To investigate the role of membrane fluidity in biofilm formation, AR3110 were treated with subinhibitory concentrations of cerulenin in both rugose and pellicle biofilm models (Supplemental material, Fig. S1). This treatment reduced membrane fluidity (t-test, P=0.0144), to levels similar those observed in the Δ*fabF* strain (Fig. 1C). The macrocolony of AR3110 typically exhibits a skin-like consistency, which prevents full resuspension of the biomass in solution (Supplemental material, Fig. S2). However, both the Δ*fabF* strain and cerulenin-treated wild-type macrocolonies acquired a buttery consistency, allowing for easy resuspension (Supplemental material, Fig. S2). The data shown in Fig. 1A and 1B demonstrated the absence of rugose macrocolony architecture and pellicle formation in both the Δ*fabF* strain and cerulenin-treated wild-type AR3110, suggesting that reduced membrane fluidity hinders the development of stable biofilm structures.

The ectopic expression of *FabB* counteracted the fluidity reduction of AR3110 caused by cerulenin treatment (*t*-test, *P*=0.8829) (Fig. 1F), thereby restoring pellicle formation in cerulenin-treated AR3110 (Fig. 1D). This observation suggests that a reduction in membrane fluidity antagonises biofilm development. While this manuscript was being prepared, a similar observation was reported in Klebsiella pneumoniae ^20^. FabR negatively regulates the expression of FabA and FabB in known Enterobacteriaceae species, including *K. pneumoniae*. Consequently, the activity of UFA synthesis in the *K. pneumoniae* Δ*fabR* mutant is elevated ^20^, leading to higher membrane fluidity compared to the WT during biofilm growth. In agreement to our observation, *K. pneumoniae* Δ*fabR* mutant was found to produce robust biofilm capable of withstanding mechanical disruptions ^20^. Dramé et al. ^20^ and this work collectively suggest that high membrane fluidity is beneficial for the robust development of microbial biofilms.

An intriguing aspect of biofilm lipid composition is the observed higher SFA content in biofilms compared to planktonic cells in several species, including *S. enterica* ^3^. This observation seems to contradict the idea that increased membrane fluidity is essential for biofilm formation. However, we hypothesize that this paradox arises from the complex relationship between lipid composition and biofilm formation. While membrane fluidity is crucial, an overabundance of UFAs may lead to excessive fluidity, compromising biofilm robustness.

In this work, we ectopically overexpressed *FabB* to enhance cis-vaccenic acid production and suppress the biofilm-inhibitory activity of cerulenin ^13^ (Fig. 1D). Strikingly, in the absence of cerulenin treatment, AR3110 cells ectopically expressing *FabB* exhibited a significantly more fluid membrane (t-test, P=0.0021) (Fig. 1F), and a pellicle was no longer formed in liquid culture (Fig. 1E). Overall, our observations demonstrated that both low and high extremes of lipid fluidity restrict biofilms from developing.

While observations across multiple bacterial species by different research groups may seem convoluted, we argue that the optimal “Goldilocks” range of membrane fluidity, required for biofilm growth versus that for planktonic growth, may vary across species. In this context, the UFA/SFA ratio and profile of phosphate headgroups can be fine-tuned to achieve the “Goldilocks” fluidity range, which we propose is instrumental in facilitating the transition from a planktonic state to a biofilm lifestyle ^3,4,20^. However, determining the precise calibration required by critical biofilm-forming bacteria for transitioning between the two lifestyles, and accounting for inter-and intra-species variations, requires extensive future research. Overall, our findings underscore the potential of targeting lipid remodeling as a strategy for biofilm control in clinical and industrial applications.

## Acknowledgement

This work is funded by an Early Career Researcher Scheme Grant from the Institute of Health and Biomedical Innovations, and an Early Career Researcher Scheme Grant from the Faculty of Health, Queensland University of Technology (Australia) to YH. YH and MT were supported by the Max Planck Queensland Centre on the Materials Science of Extracellular Matrices. The Ian Potter Foundation sponsored the CLARIOStar high-performance microplate reader (BMG, Australia). The funders had no role in study design, data collection and analysis, decision to publish, or preparation of the manuscript. The AR3110 and AR3110 ΔcsgBA strains used in this work was generously shared by Dr Cécile Bidan (Max Planck Institute of Colloids and Interfaces, Germany).

## Author contributions

YH conceptualised the project, designed and conducted the experiments, and contributed to data collection and analysis; JQ contributed to experimental resources; YH and MT supervised the study and obtained the funding. YH wrote the manuscript, and all authors edited the manuscript.

## Ethics statement

This study did not involve any human participants or animal subjects.

## Data availability

All data generated or analyzed in this study were included in this article.

