## Supplemental material for "Not too rigid or too wobbly: defining an optimal membrane fluidity range essential for biofilm formation in *Escherichia coli*"

### Materials and Methods

**Bacterial strains, plasmids, and growth conditions.** The strains used in this study were described in manuscript. The plasmid pFabB was made by PCR amplifying the *E. coli fabB* gene using the primer pair, EcfabB_Fw (CATCCGGATCCATGAAACGTGCAGTGATTAC) and EcfabB_Rv (GCTTACTGCAGTTAATCTTTCAGCTTGCGCA), then cloned into pSU2718. Growth media used in this work include: YESCA agar (1 g $L^{-1}$ yeast extract, 10 g $L^{-1}$ casmino acids, and 20 g $L^{-1}$ bacteriological agar) and LB (lysogeny broth) Lennox agar (5 g $L^{-1}$ yeast extract, 5 g $L^{-1}$ NaCl, 10 g $L^{-1}$ tryptone, and 15 g $L^{-1}$ bacteriological agar). LBON is LB without the addition of NaCl. For cases where broth culture is used, bacteriological agar is removed from the medium formulation. Wherever appropriate, antibiotics were added at the following concentrations: ampicillin (100 µg $mL^{-1}$), kanamycin (50 µg $mL^{-1}$), and chloramphenicol (17 µg $mL^{-1}$).

**Strain and plasmid constructions.** Allelic exchange mutations in this work were constructed using a modified version of the Datsenko and Wanner method [1,2], with previously described *fabF* KO primers [3]. Polyethene-glycol-induced transformations were used to transfer plasmids into the testing strains constructed for this work [4].

**Rugose macrocolonies morphology and imaging.** For rugose biofilm development, overnight bacterial cultures were first normalised to $OD_{600}$ 1.0, and the culture samples (1 µL) were spotted onto YESCA agar (10 g/L casamino acids, 1g/L yeast extract, 20 g/L Gibco™ Bacteriological agar) supplemented with 20 µg/mL Coomassie Blue. Bacteria were incubated at 26 °C for 72 hrs unless otherwise indicated. Wherever cerulenin (stock at 5 mg/mL dissolved in 200-proof ethanol) is added to YESCA agar, carrier control (ethanol) would also be included in the assessment. Sample images were taken by RedMi Note 12 Pro+ and supported with APEXEL Professional Macro Photography Lens (Item model number: HB100MM).

**Pellicle biofilm growth and staining.** Overnight bacterial cultures grown in salt-free LB were first normalized to $OD_{600}$ 1.0 and inoculated onto 1 mL liquid media in NUNC 48 plate and left to grow at 26 °C statically for 72 hrs. Culture suspensions were carefully removed, and the well was washed twice with 1x PBS. Each well was then stained with 1 mL 0.1% w/v crystal violet (dissolved in $ddH_2O$) for five mins, and then again washed twice with 1x PBS. 1 mL 1x PBS was then added to the wells to float the stained pellicle, and images were taken by ChemiDoc™ MP Imaging system using the Bio-Rad white sample tray (Coomassie Blue Stain setting). Whenever applicable, cerulenin and IPTG were added to the sample wells during inoculations.

**Membrane fluidity assay.** Overnight bacterial cultures were diluted to $OD_{600}$ 1.0, then 10 µL samples were spotted onto YESCA agar and grown overnight for ~ 30 hrs at 26 °C. Cells were then harvested from the agar surface, adjusted to $OD_{600}$ 0.5, and washed twice with 1x PBS. PMBN (dissolved in $ddH_2O$ at 5 mM) was then added to the samples at 5 µM and aerated at 26 °C for 5 mins to permeabilise the outer membrane. 1,6-diphenyl-1,3,5-hexatriene (DPH – dissolved in tetrahydrofuran to 1 mM) was added to the samples at 4 µM. The samples were then incubated at 26 °C for 40 min. Unincorporated DPH was removed by two washes with 1x PBS and then resuspended in PBS. The fluorescence anisotropies (*r*) of the samples (200 µL volume), which negatively correlates to membrane fluidity, was calculated as previously described [5].

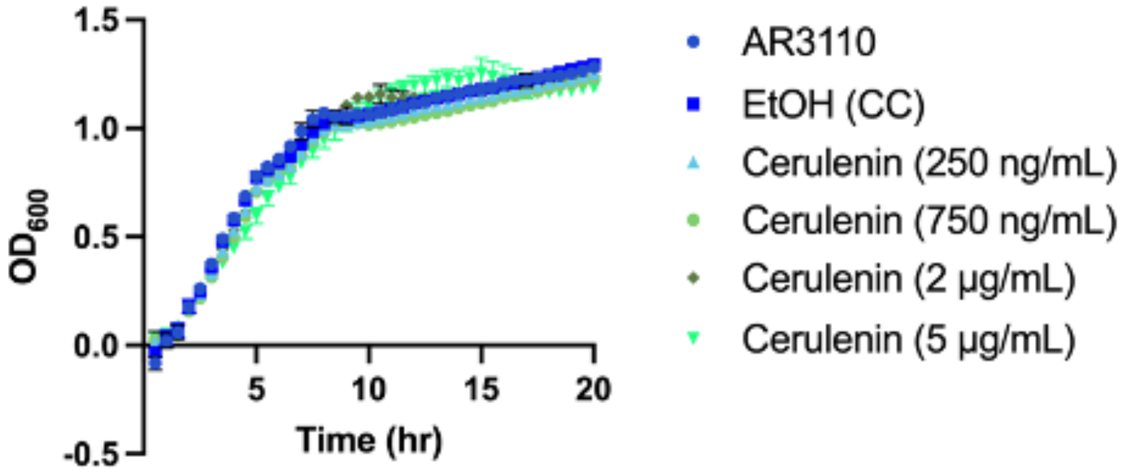


**Fig S1.** AR3110 growth in the presence of cerulenin.

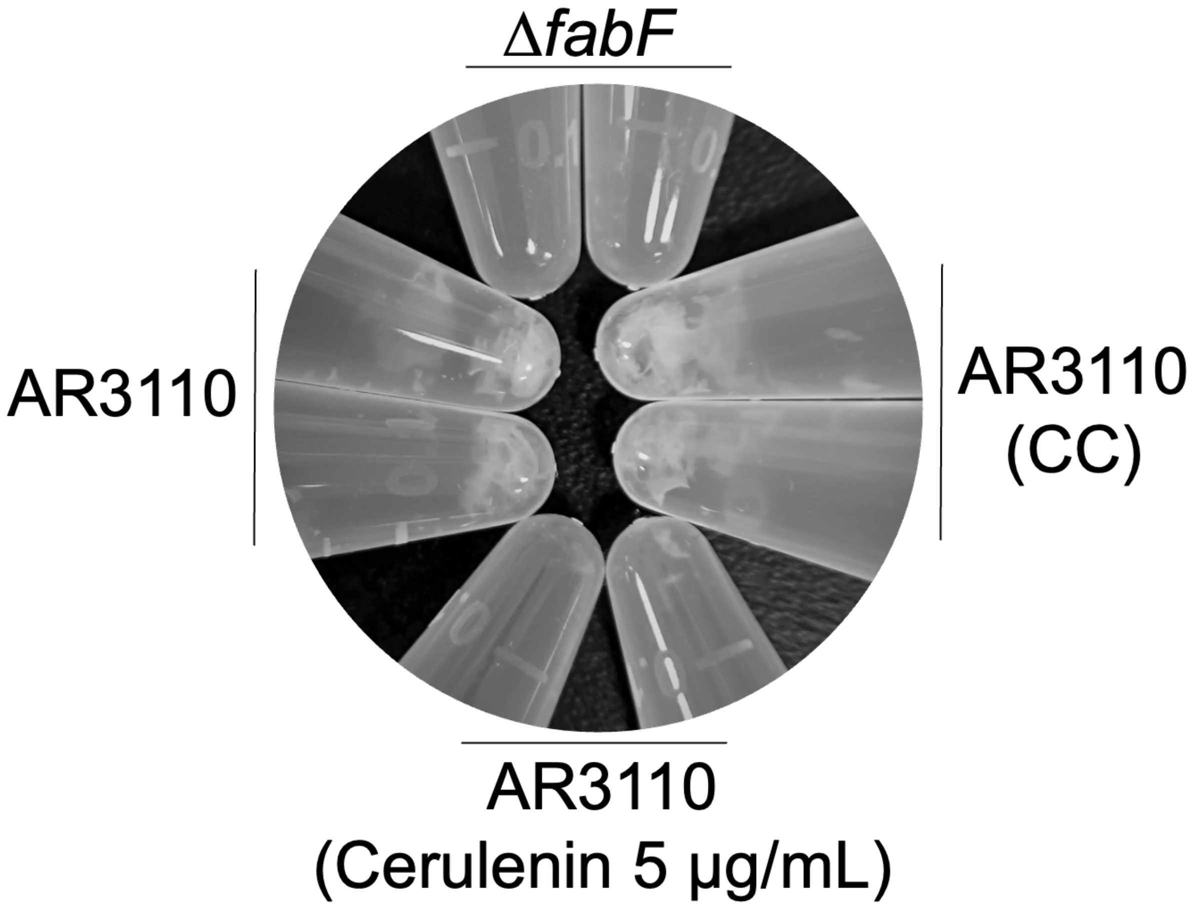


**Fig S2.** The *fabF* null mutation and cerulenin treatment attenuated the skin-like consistency of AR3110 biofilms. Overnight cultures were inoculated onto YESCA agar as per the description for developing rugose biofilm. On day 3, the entire macrocolonies were scrapped from the agar surface with a 10 μL plastic loop and resuspended in 1× PBS. The pellet was resuspended repeatedly with 1000 μL pipette for 15 sec and then vortexed at max capacity for 20 sec. Both *fabF* null mutant and cerulenin treated macrocolonies were readily resuspended, while the AR3110 untreated controls retained skin-like meshes in solution. The figure imaged is representative of three independent repeats.
